# Inversion Genotyping in the *Anopheles gambiae* Complex Using High-Throughput Array and Sequencing Platforms

**DOI:** 10.1101/2020.05.25.114793

**Authors:** R. Rebecca Love, Marco Pombi, Moussa W. Guelbeogo, Nathan R. Campbell, Melissa T. Stephens, Roch K. Dabire, Carlo Costantini, Alessandra della Torre, Nora J. Besansky

## Abstract

Chromosomal inversion polymorphisms have special importance in the *Anopheles gambiae* complex of malaria vector mosquitoes, due to their role in local adaptation and range expansion. The study of inversions in natural populations is reliant on polytene chromosome analysis by expert cytogeneticists, a process that is limited by the rarity of trained specialists, low throughput, and restrictive sampling requirements. To overcome this barrier, we ascertained tag single nucleotide polymorphisms (SNPs) that are highly correlated with inversion status (inverted or standard orientation). We compared the performance of the tag SNPs using two alternative high throughput molecular genotyping approaches versus traditional cytogenetic karyotyping of the same 960 individual *An. gambiae* and *An. coluzzii* mosquitoes sampled from Burkina Faso, West Africa. We show that both molecular approaches yield comparable results, and that either one performs as well or better than cytogenetics in terms of genotyping accuracy. Given the ability of molecular genotyping approaches to be conducted at scale and at relatively low cost without restriction on mosquito sex or developmental stage, molecular genotyping via tag SNPs has the potential to revitalize research into the role of chromosomal inversions in the behavior and ongoing adaptation of *An. gambiae* and *An. coluzzii* to environmental heterogeneities.

## INTRODUCTION

A chromosomal inversion is a structural mutation that arises when a chromosome segment breaks and reattaches in reverse orientation. Those that are retained as long-term polymorphisms often span hundreds or thousands of genes (Wellenreuther & Bernatchez 2018). Suppressed recombination in inversion heterozygotes (between inverted and non-inverted orientations) preserves allelic combinations on the inverted arrangement as haplotype blocks. The proposed role of inversions in adaptation to environmental heterogeneities arises from the expectation that locally adapted haplotype blocks can be maintained by spatially and temporally varying selection (Hoffmann & Rieseberg 2008; Kirkpatrick 2010; Kirkpatrick & Barton 2006; Schaeffer 2008). Despite significant advances in genomic resources and technologies, a more detailed understanding of the precise alleles targeted by selection and their phenotypic consequences is largely lacking. One important roadblock to progress is the paucity of methodologies that allow inversion genotyping at scale.

Polymorphic chromosomal inversions are abundant in all four of the major malaria-transmitting mosquitoes with pan-African distributions (Ayala *et al.* 2017; Coluzzi *et al.* 2002), including the two sister species of the *Anopheles gambiae* complex studied here, *An. gambiae sensu stricto* (hereafter *An. gambiae*) and *An. coluzzii*. It has been suggested that these inversion polymorphisms promote ecological flexibility, enabling the successful exploitation of heterogeneous environments across tropical Africa (Ayala *et al.* 2017; Coluzzi *et al.* 2002; Costantini *et al.* 2009). Decades ago, intensive and laborious cytogenetic studies demonstrated that inversion frequencies correlate with latitudinal gradients of aridity (Coluzzi *et al.* 1979), seasonal fluctuations in rainfall (Rishikesh *et al.* 1985), and local microhabitat differences (Coluzzi *et al.* 1979), patterns that persist stably over decades and across different geographic regions. These observations suggest that inversions confer an adaptive benefit in arid environments, where they achieve their highest frequencies. The epidemiological relevance of inversion polymorphism for malaria transmission and control, beyond habitat expansion and seasonal persistence of the mosquito disease vectors, was manifest by differences in vector house resting behavior (Coluzzi *et al.* 1979; Molineaux & Gramiccia 1980). The significantly greater tendency of the inversion-carrying fraction of the mosquito population to rest indoors caused non-uniform exposure to indoor-based vector control, reducing its efficacy.

Despite the undeniable public health importance of this phenomenon, scientific understanding has barely advanced in the forty years since its initial discovery. No small reason for this hiatus is the technological and logistical difficulty of inversion genotyping. Polytene chromosome analysis of anopheline mosquitoes (della Torre 1997) is the current basis of inversion genotyping. Strongly rate-limiting, sex-specific and stage-specific, it requires dissection, preparation, and microscopic analysis of ovarian polytene chromosomes by expert cytogeneticists with highly specialized training in the interpretation of chromosome banding patterns of the focal species.

DNA-based molecular assays would offer a much more rapid and widely available approach to inversion genotyping. These could be applied to mosquitoes regardless of sex, developmental stage, or method of preservation, and would require no more training than that normally associated with any molecular entomology laboratory. More than ten years ago, a rapid PCR assay was developed for genotyping of the 22-Mb 2La inversion (White *et al.* 2007), one of six common inversion polymorphisms in *An. gambiae* (all shared with *An. coluzzii* except one, 2Rj). The availability of this assay, which targets 2La inversion breakpoints, simplified the search for phenotypic traits and single nucleotide polymorphisms (SNPs) associated with the inverted orientation (2La). Studies employing this tool in the laboratory suggested that the inverted orientation was associated with increased thermal and desiccation tolerance, a thicker cuticle, higher body water content, a more aggressive upregulation of heat-responsive genes such as heat shock genes, and a higher energy budget relative to the alternative (2L+^a^) arrangement (Cassone *et al.* 2011; Cheng *et al.* 2018; Fouet *et al.* 2012; Gray *et al.* 2009; Reidenbach *et al.* 2014; Rocca *et al.* 2009). Importantly, this inversion genotyping tool also was used to advance the study of natural populations. Inversion 2La was found to be associated with a reduced tendency to rest indoors and a lower malaria oocyst infection prevalence, corroborating historical evidence based on cytogenetic analysis (Coluzzi *et al.* 1979; Petrarca & Beier 1992; Riehle *et al.* 2017). Association mapping employing population pools of alternative 2La homokaryotypes revealed dozens of candidate SNPs significantly associated with desiccation tolerance (Ayala *et al.* 2019).

However, molecular genotyping tools that perform robustly for the common inversions on the right arm of chromosome 2 (2Rj, 2Rb, 2Rc, 2Rd, 2Ru) have been lacking, with the sole exception of a newly available set of polymerase chain reaction (PCR) restriction fragment length polymorphism (RFLP) assays for 2Rb (Montanez-Gonzalez *et al.* 2020). Two previously developed PCR genotyping assays, one that targeted the breakpoints of this inversion and another that targets the breakpoints of inversion 2Rj, proved unreliable or had limited geographic application in natural populations, presumably due to structural variation in inversion breakpoint regions (Coulibaly *et al.* 2007; Lobo *et al.* 2010). The newly developed PCR-RFLP genotyping assays for 2Rb (Montanez-Gonzalez *et al.* 2020), and additional genotyping assays under development for 2Rc (Montanez-Gonzalez, Vallera, Calzetta, Love, Pombi, Guelbeogo, Dabire, Costantini, Pichler, Petrarca, della Torre, Besansky, unpublished) exploit tag SNPs inside the rearranged region whose allelic state is strongly correlated with inversion orientation regardless of their position relative to the breakpoints (Love *et al.* 2019). To our knowledge, no DNA-based molecular assays exist for the genotyping of the other inversions in *An. gambiae* or *An. coluzzii*.

We recently described a strategy that exploited the *An. gambiae* and *An. coluzzii* database of natural variation (Ag1000G; www.malariagen.net/projects/ag1000g) (Miles *et al.* 2017) to identify tag SNPs predictive of inversion orientation for all six common inversion polymorphisms in these species. Using these tags, we developed an algorithm capable of *in silico* inversion genotyping based on SNPs called from whole genome resequencing data (Love *et al.* 2019). This is a rapid and powerful approach assuming that whole genome sequence data are already available or will be produced for other reasons. However, it does not satisfy experimental designs in which genomic sequence data is not otherwise required, and where its procurement would be cost-prohibitive. For the requisite statistical power, studies aimed at finding significant associations between inversions and behavioral or physiological phenotypes will likely require thousands of specimens of known inversion genotype. Here, we develop cost-effective high-throughput molecular methods of inversion genotyping to address this need. Using tag SNPs ascertained in Ag1000G (Love *et al.* 2019), we compare two molecular platforms that allow inversion genotyping of hundreds or thousands of individual *An. coluzzii* and *An. gambiae* mosquitoes at tens or hundreds of tag SNPs targeting all six inversions in a single experiment. One platform, the TaqMan OpenArray (Life Technologies), referred to hereafter as OA, is a 5’-exonuclease method that genotypes tag SNPs based on PCR in the presence of allele-specific probes, both labeled with different reporter dyes. The other, Genotyping-in-Thousands by sequencing (GT-seq), is a custom amplicon sequencing approach that genotypes tag SNPs by next-generation sequencing of multiplexed PCR products (Campbell *et al.* 2015). Using 960 individual *An. gambiae* and *An. coluzzii* mosquitoes previously karyotyped cytogenetically and up to 184 SNP markers, we show that both approaches successfully predict inversion genotypes for the common polymorphic inversions in these species (excluding 2Rc in *An. gambiae* and 2Rd in *An. coluzzii*). Our data suggest that these methods not only offer efficiency of scale and cost, but also represent substantial improvements in genotyping accuracy relative to the classical cytogenetic approach.

## MATERIALS AND METHODS

### Mosquito study population

Burkina Faso lies in the arid Sudan savanna belt of West Africa. In this region, *An. gambiae* and *An. coluzzii* are highly polymorphic for chromosomal inversions (Costantini *et al.* 2009). Sampling was conducted in a 35 × 65 km area located 30 km SW of the capital, Ouagadougou. In total, 85 villages approximately 5 km distant from each other were sampled in 2006. Mosquito collection was performed indoors in the early afternoon by pyrethrum spray catch in 3-5 compounds per village. Morphological identification and initial processing was performed in the field under a dissecting microscope. *An. gambiae s.l*. females at the appropriate stage for polytene chromosome analysis were each assigned a unique numerical code, whose value was incremented by ‘1’ with each new mosquito as the collection progressed. Ovaries of each female mosquito were immediately cropped and placed in an individual 1.5 ml tube containing Carnoy’s fixative (1:3 glacial acetic acid:absolute ethanol), labeled with its unique numerical code. The corresponding carcass was placed in an individual 1.5 ml tube containing a desiccant (silica gel) and a matching numeric label unique for that mosquito. Ovaries were stored at −20°C, and carcasses maintained at ambient temperature before further processing.

Mosquito DNA was extracted from the carcass using a CTAB method (Chen *et al.* 2010) and identified to species using rDNA-based PCR assays (Favia *et al.* 1997; Scott *et al.* 1993). The corresponding ovaries were prepared for karyotype analysis according to standard procedures (della Torre 1997). The banding pattern was observed under a phase-contrast microscope (400x) and interpreted with reference to the cytogenetic map (George *et al.* 2010; Pombi *et al.* 2008). Karyotype analysis was performed on >1,770 mosquitoes. The effort was divided equally between two groups without spatial or temporal sampling bias, based strictly on mosquitoes with odd-versus even-valued numerical codes. For this study, we selected a subset of 960 mosquitoes based on their cytogenetic karyotypes, with the goal of achieving maximum possible inversion genotype balance for the purpose of validating the molecular tag SNPs.

### OA and GT-seq assay design and genotyping

Because the methods we used to develop and validate the set of tag SNPs for *in silico* inversion genotyping (Love *et al.* 2019) were still being refined at the time the present study was initiated, the initial list of candidate molecular tags differed slightly from the *in silico* set, though overlap was extensive. One minor methodological difference in ascertainment was that, for the molecular tag SNPs, genotypic concordance was not based on ten bootstrap replications of a training set, but instead was based on the simple percentage of mosquitoes with a matching inversion- and SNP-genotype out of the total mosquito sample analyzed for that inversion in the Ag1000G variation database. This was calculated separately for each of three inversion genotypes [homozygous standard (*i.e.*, uninverted), heterozygous, homozygous inverted], with the minimum value taken as the conservative genotypic concordance. The other methodological difference was that ascertainment of tag SNPs in 2Rc was based on slightly different data partitions: (i) *An. coluzzii*, and (ii) *An. gambiae* after exclusion of BAMAKO (Manoukis *et al.* 2008) and specimens carrying the inverted arrangement of 2Ru. Beginning with a ranked list of tags based on descending genotypic concordance, we applied filters (described below) that narrowed the numbers of tags based on design criteria unique to each platform.

#### *OA:* TaqMan assays were designed by the Dana-Farber/Harvard Cancer

Center (DF/HCC) Genotyping and Genetics for Population Sciences Core, a unit of the Partners HealthCare Center for Personalized Genetic Medicine. Assays designed for this platform require forward and reverse PCR primers which produce ~100 base amplicons containing the tag SNP, and additionally require two allele-specific fluorescently labeled 30-bp probes (‘reporters’) that discriminate between the reference allele at the tag (VIC dye) and the alternate allele (FAM dye). Candidate SNPs were filtered out if they were surrounded by runs of nucleotides and low complexity regions that interfered with acceptable primer design parameters. Further filtering was performed if candidate SNPs were surrounded by high frequency variants in the 25 bases immediately upstream or downstream of the tag SNP. High frequency was defined as ≥5% in at least one inversion genotype (*i.e.*, homozygous standard, heterozygous, homozygous inverted) or at least two population samples analyzed for a given inversion in the Ag1000G database. [As discussed in Love *et al*. (2019), the population samples analyzed varied depending upon the inversion under consideration, due to inferred taxonomic or geographic population structure based on principal components analysis]. Such high frequency variants were deemed likely to significantly interfere with successful probe annealing. Due to the highly polymorphic nature of the *An. gambiae* genome (Miles *et al.* 2017), these filters eliminated many candidate tags. To ensure that we retained at least six tag SNPs per inversion for genotyping, we were compelled to reduce the genotypic concordance threshold below the 0.8 level imposed in Love *et al*. (2019). Even lowering the threshold to 0.7 for 2Rc tags in *An. gambiae* failed to yield more than three candidates, and we declined to reduce that threshold further. Minimum genotypic concordances for each inversion were 0.7 for 2Rb, 2Rc and 2Ru; 0.75 for 2Rd; 0.9 for 2Rj; and 0.9925 for 2La.

After filtering, we retained 54 tag SNPs in total, ranging from 6 to 11 per inversion except 2Rc in *An. gambiae,* with only 3 tag SNPs. Based on these 54 tags, we selected a custom 64-assay TaqMan OpenArray genotyping plate design whose 3,072 reaction through-holes are divided into 48 sub-arrays, each with 64 through-holes (54 of which were preloaded with a single custom assay). One such plate genotypes 48 mosquitoes at 54 tags (2,592 genotypic assays).

DNA quantification of genomic DNA from 960 mosquitoes was conducted by DF/HCC via picogreen-based fluorimetry; average DNA concentration was 26 ng/ul (range, 0.1-58.1 ng/ul). OA requires 250 copies of a haploid genome for each individual through-hole [0.0675 ng of *An. gambiae* genomic DNA, assuming a haploid genome size of 0.27 pg (260 Mb); (Besansky & Powell 1992)]; 64 through-holes require only ~4-5 ng DNA per mosquito. DF/HCC performed the genotyping using endpoint detection of fluorescent signals on the TaqMan OpenArray Genotyping System, following manufacturer’s specifications (Applied Biosystems, Foster City CA, USA). Conditions for genotyping are available upon request to DF/HCC. Tag SNPs, primers and probes for genotyping assays are provided in Table S1.

*GT-seq:* GTseek LLC conducted multiplex primer design and consulted on GT-seq optimization. Because this multiplexed amplicon sequencing approach uses only unlabeled PCR primers to produce 50-100 bp amplicons spanning a tag SNP, high frequency variants neighboring the tag are not a limitation. However, the highly multiplexed nature of GT-seq, allowing simultaneous amplification of up to 500 SNP loci per individual for thousands of individuals, requires that the primer pool be optimized not only for individual amplicons (*e.g*., by avoiding nucleotide runs, low complexity regions, and primer-dimer), but also to minimize primer interactions across loci and mis-priming with other amplicons. The initial list of candidate tag SNPs ranked by concordance was filtered based on the output of custom perl scripts to evaluate primer pools (Campbell *et al.* 2015, https://github.com/GTseq). Minimum concordance values varied by inversion (>0.8 for 2Rc; >0.85 for 2Rd; >0.9 for 2Rb, 2Rj, and 2Ru; >0.995 for 2La). Candidate tag SNPs for 2La, which were overly abundant, were pruned by selecting every third candidate from a list ordered by chromosome position.

Following Campbell *et al.* (2015), Illumina sequencing primer sites were added to locus-specific forward and reverse primer sequences to create PCR1 primers, which were ordered along with PCR2 primers (a set of 96 i5 and i7 indexes) from Integrated DNA technologies (IDT) in 96-well plate format at a 25nmole synthesis scale and a concentration of 200 μM in Tris-EDTA pH 8.0 buffer. GT-seq test libraries were prepared and sequenced by the University of Notre Dame Genomics and Bioinformatics Core Facility (GBCF) from a subset of specimens (n=192) to refine preparation techniques and identify primers that produced PCR artefacts or were overrepresented. Following optimization, primer pools were re-made to include only the optimized panel of PCR 1 primers. Tag SNPs and PCR 1 primers for GT-seq genotyping are listed in Table S2.

The final libraries prepared by the GBCF included the same 192 specimens used during optimization and 765 additional specimens. They were constructed without optional exo-SAP treatment following Campbell *et al.* (2015), with the following modifications to PCR conditions and post library cleanup: PCR1: 95 °C – 15 min; 5 cycles [95 °C – 30s, 3% ramp down to 57°C-30s, 72°C-2min]; 10 cycles [95 °C-30 s, 65 °C-30 s, 72 °C-30 s]; 4 °C hold. PCR2: 95 °C - 15 min; 10 cycles [95°C-10s; 62°C-30s; 72°C-30s]; 72°C-5min; 4°C hold. Following PCR2, each plate of samples was purified and normalized using the Just-a-Plate 96 PCR Purification and Normalization Kit (Charm Biotech) according to manufacturer’s instructions. Following normalization, 10 ul of each sample per 96 well plate (up to 960 ul total) was then combined into a 1.5-mL Eppendorf tube, for a total of 10 tubes. From each tube, 300ul was transferred to a fresh 1.5-mL Eppendorf tube for two rounds of purification using AMPure XP paramagnetic beads (Beckman Coulter, Inc.) with ratios of 0.5X and 1.3X respectively. Purified libraries were eluted in 35 ul 1xTE and transferred to fresh 1.5-mL tubes before adding 3.5 ul buffer EB containing a 1% Tween 20 solution.

Each of the 10 plate libraries was quality assessed on an Agilent Bioanalyzer 2100 High Sensitivity chip and quantified by qPCR using the Illumina Kapa Library Quantification Kit (Roche, Cat. #KK4824). The libraries were then normalized to a concentration of 4 nM and pooled for sequencing. The final pooled library containing 957 *An. gambiae* and *An. coluzzii* individuals was sequenced on a single lane of Illumina NextSeq 500 v2.5 (75 cycle) High Output flowcell using a dual indexed 75bp single-end read. Base calling was done by Illumina Real Time Analysis (RTA) v2 software.

Using scripts described in the bioinformatics pipeline of Campbell *et al.* (2015) and available on Github (https://github.com/GTseq), sequencing data were demultiplexed into single fastq files for each individual sample. Individuals were genotyped at each locus with a perl script (GTseq_Genotyper_v3.pl) that counts the occurrence of each allele at a locus within individual fastq files. The ratio of allele 1 to allele 2 counts was used to generate a genotype for each locus with total read counts >10, following the methods and cut-offs of Campbell *et al.* (2015).

### Filtering and calling multilocus inversion genotypes

The procedures for filtering and calling molecular inversion genotypes were the same for both OA and GT-seq platforms. Filtering steps were as follows. For each tag SNP, we calculated the percentage of mosquito specimens in the sample with a genotype call at that tag (the SNP call rate). If SNP call rates were <80%, the underperforming tag SNPs were eliminated from further analysis. In addition, for each mosquito specimen analyzed, we calculated the percentage of tag SNPs with a genotype call (the specimen call rate). If the specimen call rate was <80%, that specimen was excluded from further analysis. Note that the mosquito specimens in the sample varied according to the inversion under consideration: 2La, 2Rb and 2Ru tags perform in both species, 2Rj and 2Rd tags are *An. gambiae*-specific, and defined subsets of 2Rc tags (referred to in this work as 2Rc_col and 2Rc_gam) apply respectively to *An. coluzzii* or *An. gambiae* individuals.

To calculate the multilocus inversion genotype for each specimen, we converted the raw genotype data for individual tag SNPs to the count of alternate alleles (if necessary), where ‘0’ is a homozygote for the reference allele, ‘1’ is a heterozygote carrying one reference allele, and ‘2’ is a homozygote for the alternate allele. Next, we averaged the number of alternate alleles present across all tag SNPs in a given inversion, and binned this average to produce a predicted inversion genotype (0-0.67, 0; 0.68-1.33, 1; 1.34-2, 2). Multilocus molecular genotypes were then compared to each other and with cytogenetically determined inversion genotypes.

### Code and data availability

Supplemental files are available at FigShare. Table S1 contains OA tag SNP ID numbers, locations, reference and alternate alleles, forward and reverse primer sequences, and probe sequences. Table S2 contains GT-seq tag SNP ID numbers, locations, reference and alternate alleles, and forward and reverse primer sequences. Table S3 contains specimen ID numbers and inversion genotypes (cytogenetic, OA, and GT-seq) for each individual mosquito. Code used to generate the data can be found on Github (https://github.com/GTseq and https://github.com/rrlove/molec_karyo_notebooks).

## RESULTS AND DISCUSSION

### OA

Custom OA plates were used to genotype 960 individual *An. gambiae* and *An. coluzzii* mosquitoes at 54 tag SNP loci. The SNP call rate for one of the 54 loci fell below the 80% threshold (77.7%) and was eliminated from the panel. After filtering, the SNP call rate averaged 99.4% for the remaining 53 tags (range, 98.3%-100%). Three specimens were dropped from analysis due to the belated determination that their cytogenetic genotypes were ambiguous. Four additional mosquito specimens were dropped from further OA analysis due to unacceptably low specimen call rates (ranging from 17.5% to 52.6%). The remaining 953 specimens had an average specimen call rate of 99.3% (range, 87.7%-100%). The final number of OA tags per each inversion, and their approximate genomic position within the inversion, are shown in Table 1 and Figure 1.

**Table 1.**
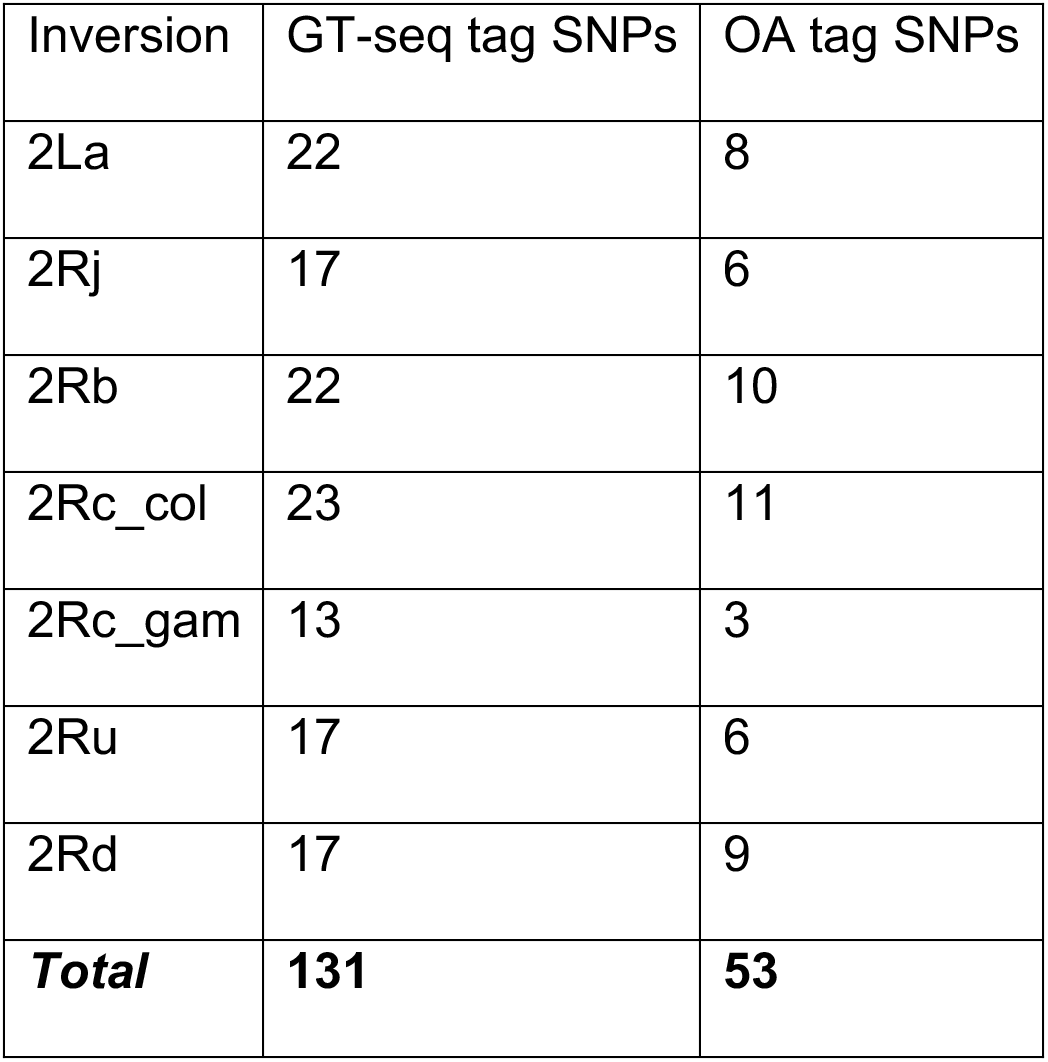
The number of tag SNPs by inversion and molecular method.

**Figure 1.**
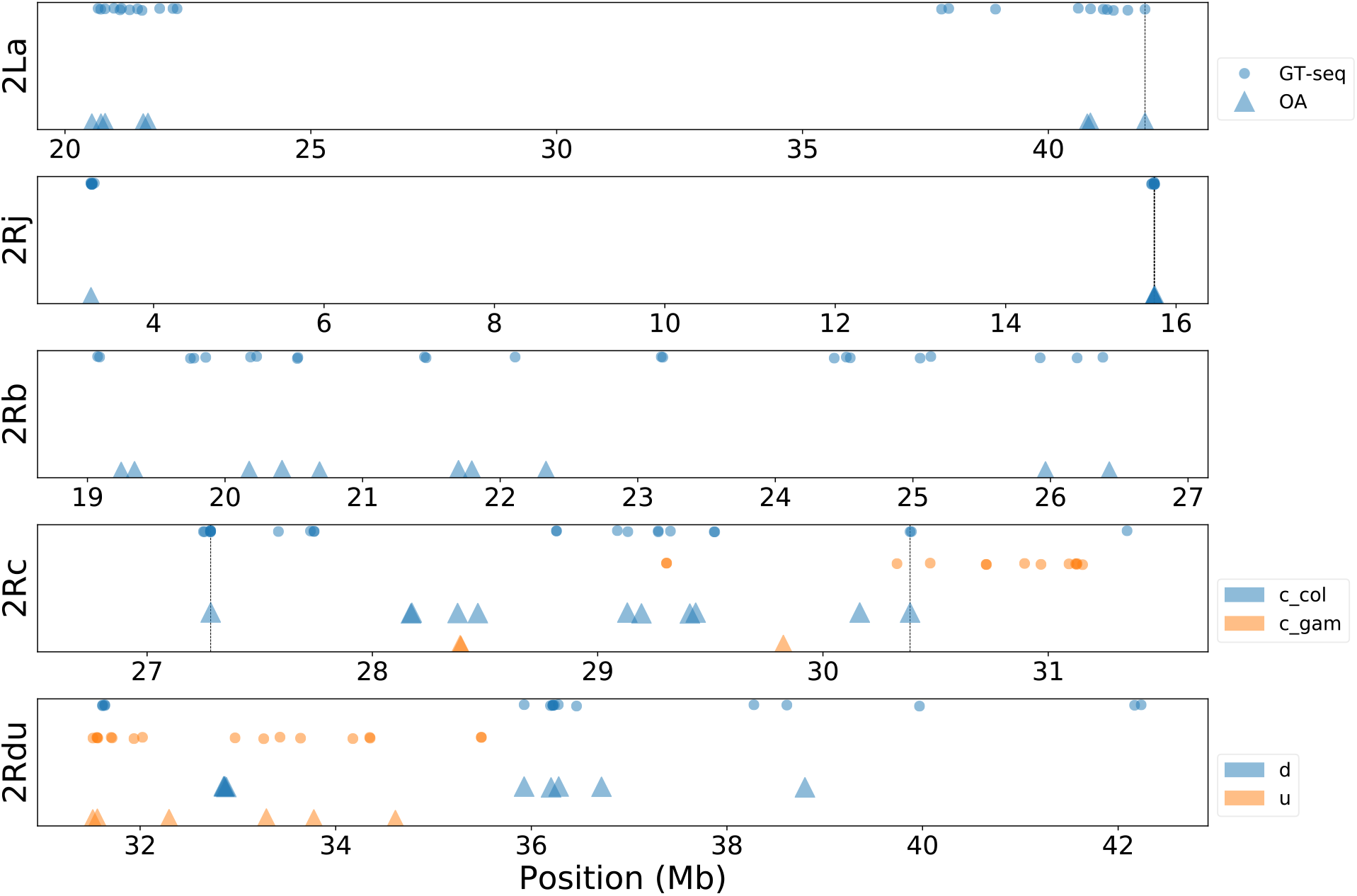
Locations of tag SNPs assayed for each inversion for OA and GT-seq. The bottom panel labeled 2Rdu shows inversions 2Ru and 2Rd on the same panel, as 2Ru is wholly encompassed by 2Rd. Vertical lines indicate SNPs common to both methods.

### GT-seq

Sequencing from one NextSeq lane included the pooled GT-seq library of 957 *An. gambiae* and *An. coluzzii,* as well as another GT-seq pooled library of 235 *An. funestus* mosquitoes pertaining to an independent experiment to be described elsewhere. This produced ~359M total reads, of which ~236M could be assigned to the 957 *An. gambiae* and *An. coluzzii* specimens based on their barcode sequences. Read counts from each of the ten *An. gambiae*-*An. coluzzii* sample plates ranged from 17.5M to 27.6M reads per plate and read counts per individual mosquito averaged 246,925 (SD 85,614). The tag SNP call rate was below the 80% threshold for three tags, which were subsequently dropped from the genotyping panel. For the remaining 131 tags, SNP call rates averaged 98.2% (range, 80.25% to 100%). As was the case for OA analysis, three specimens were dropped due to ambiguous cytogenetic genotypes. Of the remaining 954, two were considered to have failed because they had specimen call rates below the 80% threshold, and were thus dropped. Those 952 within acceptable limits had average specimen call rates of 98.2% (range, 93.1% to 100%). The final number of GT-seq tags per each inversion, and their approximate genomic position within the inversion, are shown in Table 1 and Figure 1.

### Concordance

The inversion genotypes inferred for each specimen by the three methods (cytogenetics, OA, and GT-seq) are provided in Table S3. We compared these genotypes to assess their concordance. Due to our filtering rules, not every specimen had genotype calls by both molecular methods. We focused our assessment on the subset of specimens that were successfully genotyped with all three methods (435 for *An. gambiae*, and 513 for *An. coluzzii*). As summarized in Figure 2 and Table 2, over 90% of the relevant mosquito samples had concordant genotypes for all three methods with the notable exception of *An. gambiae* genotyped for 2Rc, where three-way concordance fell to ~81%. We discuss the special case of 2Rc in *An. gambiae* more fully below; here, we concentrate on the five other inversions and 2Rc in *An. coluzzii*.

**Table 2.** Concordance of genotypes imputed by cytogenetics (CYT), OA, and GT-seq for each inversion.

|  | 2La, N (%) | 2Rj, N (%) | 2Rb, N (%) | 2Rc-col, N (%) | 2Rc-gam, N (%) | 2Rd, N (%) | 2Ru, N (%) |
| --- | --- | --- | --- | --- | --- | --- | --- |
| Concordance: |  |  |  |  |  |  |  |
| Three-way | 875 (92.30) | 434 (99.77) | 864 (91.14) | 483 (94.15) | 352 (80.92) | 420 (96.55) | 870 (91.77) |
| Discordance: |  |  |  |  |  |  |  |
| CYT vs (GT-seq + OA) | 73 (7.70) | 1 (0.23) | 41 (4.32) | 18 (3.51) | 39 (8.97) | 14 (3.22) | 56 (5.91) |
| All other | 0 (0) | 0 (0) | 43 (4.54) | 12 (2.34) | 44 (10.11) | 1 (0.23) | 22 (2.32) |
| (CYT + GT-seq) vs OA | 0 (0) | 0 (0) | 43 (4.54) | 11 (2.14) | 31 (7.13) | 1 (0.23) | 19 (1.05) |
| (CYT + OA) vs GT-seq | 0 (0) | 0 (0) | 0 (0) | 1 (0.19) | 10 (2.30) | 0 (0) | 3 (0.32) |
| Three-way | 0 (0) | 0 (0) | 0 (0) | 0 (0) | 3 (0.69) | 0 (0) | 0 (0) |
| Specimens assayed by all 3 methods | **948** | **435** | **948** | **513** | **435** | **435** | **948** |

**Figure 2.**
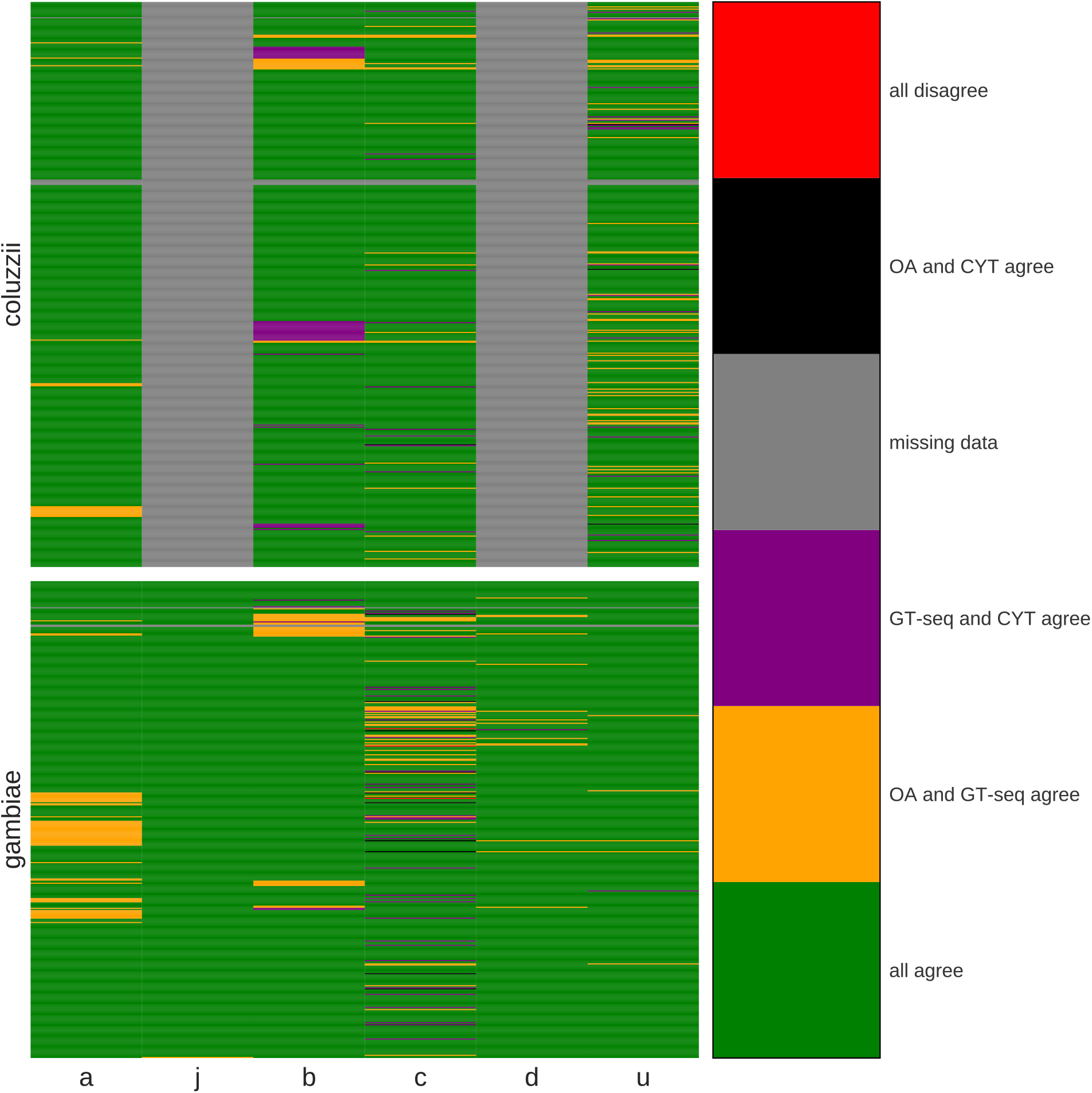
Concordance heat map of genotypes imputed by cytogenetics (CYT), OA, and GT-seq. Each row is an individual mosquito, and each column compares inversion genotypes derived from three genotyping approaches for a given inversion (a, 2La; j, 2Rj; b, 2Rb; c, 2Rc; d, 2Rd; u, 2Ru). Rows are grouped by species; 2Rj and 2Rd tags are not applicable in *An. coluzzii*. Green represents 3-way genotypic concordance; yellow, concordance between OA and GT-seq; purple, concordance between CYT and GT-seq; black, concordance between CYT and OA; gray is missing data; red is 3-way discordance.

A strikingly high number of specimens had multilocus molecular genotypes inferred from both OA and GT-seq that agreed, but were jointly discordant with cytogenetics. Except for 2Rj with negligible discordance (and correspondingly low levels of polymorphism in our sample), the cytogenetic versus multilocus molecular discordance affected from 14 to 73 mosquito specimens per inversion, representing 3% to 8% of the mosquito samples (mean, 5%). Although cytogenetic karyotyping may be considered the gold standard for inversion genotyping, two important considerations lend considerable confidence to molecular genotypes, particularly when both molecular approaches concur. First, while none of the tag SNPs are deterministic (*i.e.*, none is perfectly and invariably correlated with inversion orientation), OA and GT-seq infer genotypes based on multiple predictive tags scored per inversion, thus providing weight of numbers. Second, the final set of tags used for OA and GT-seq are almost completely non-overlapping, an outcome produced by distinct filters imposed on the initial list of candidate tags during assay development (see Methods; Figure 3). Accordingly, agreement between both molecular methods is even stronger evidence in favor of the inferred molecular genotype than that provided by one or the other molecular method by itself. Table 2 shows that the two molecular methods agree at least 95% of the time (an average of 98%), except in the case of 2Rc in *An. gambiae*.

**Figure 3.**
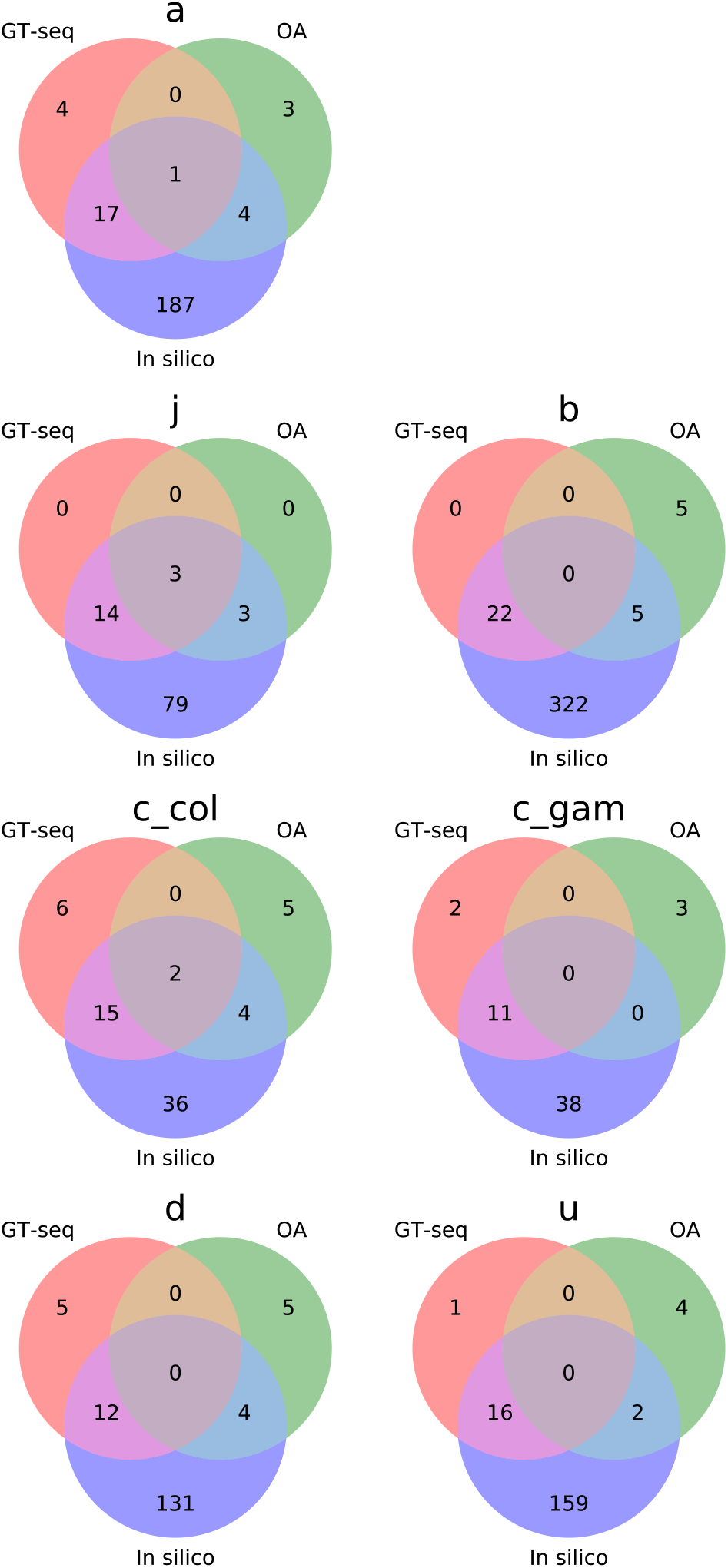
Venn diagrams showing degree of overlap between tag SNPs developed for *in silico* inversion genotyping by Love *et al*. (2019) and those developed in this study for OA and GT-seq.

We hypothesized that some genotypic discordances, specifically those in which the two molecular methods agree but conflict with cytogenetics, are caused by cytogenetic errors rather than systematic biases in the molecular approaches. This is difficult to demonstrate conclusively, because the specimens used to compare the three genotyping methods have not been subjected to whole genome sequencing. Furthermore, it was not possible to double-check the cytogenetic karyotypes of the specimens with discordances in the majority of cases, because neither slides nor ovaries were available. However, there is strong evidence consistent with cytogenetic error. During sampling in the field, specimens were assigned numerical identifiers that incremented by “1” throughout the process; cytogenetic karyotyping was later split between two institutions on the basis of even- or odd-numbered identifiers (see Methods). This procedure virtually eliminates the possibility of biases owing to temporal or spatial heterogeneities during the course of the mosquito sampling. Because even- and odd-numbered specimens should be random subsamples from the same populations, we would expect no difference in discordance rates between them. This was not what we found. Of the specimens assayed by both molecular methods in this study, 280 were odd-numbered and 677 even-numbered. Focusing on the inversions with the largest numbers of specimens whose cytogenetic and joint molecular genotypes disagreed (2La, 73; 2Rb, 41; 2Ru, 56; Table 2), the combined 170 such discordances occurred disproportionately in even-numbered specimens: 169 of the 170. Analyses of 2×2 contingency tables demonstrated highly significant departures from the null hypothesis (by Chi-square and Fisher exact probability tests), consistent with the notion that cytogenetic error disproportionately affecting the even-numbered specimens is responsible for these genotypic discrepancies (~5%). If this is the case, then based on the fact that both molecular approaches agree >95% of the time, we suggest that the true error rate for either molecular approach is <5%, probably closer to ~2%.

It is important to recognize that although the tag SNPs assayed by the two molecular approaches are largely non-overlapping for technical reasons, the assumptions underlying the ascertainment of the initial set of candidate tags were the same. The implication is that if those assumptions are violated in natural populations, both approaches may agree on the wrong genotype. The tags were ascertained in the Ag1000G variation database, whose content was heavily biased toward *An. gambiae* at the time of their discovery (Love *et al.* 2019). Available samples of *An. coluzzii* were more limited in numbers and geographic representation, although Burkina Faso was one of two countries represented for this species. In addition to the issue of sampling limitations is the issue of unsuspected (cryptic) population structure that could affect the performance of these tags. Population structure could arise from several non-mutually exclusive scenarios: (i) a lack or reduction of connectivity between natural populations; (ii) local heterogeneity in selection pressures acting on targets inside the inversion; and/or (iii) violation of the assumption that the focal inversion arose uniquely (*i.e.*, has a monophyletic origin). We suspect that at least one of these scenarios applies to 2Rc tags ascertained in *An. gambiae*, probably explaining their lower rate of apparent success in genotyping (based on lower concordance values across the board; Table 2). Previous work has shown that applying candidate tags to a taxon in which they are not valid has the effect of downwardly biasing the average number of inferred alternate alleles (Love *et al.* 2019). Consistent with this, if 2Rd tags in the present study were inappropriately applied to *An. coluzzii*, the vast majority (38 of 39) of specimens genotyped cytogenetically as heterozygotes would be molecularly genotyped as standard homozygotes (Table S3). Based on this, we expect the systematic underestimation of the number of alternate alleles to produce a distinctive pattern where ‘true’ standard homozygotes are correctly identified, but heterozygotes and inverted homozygotes would be incorrectly genotyped molecularly as standard homozygotes. Table 3 shows the distribution of discordant genotypes between cytogenetics and joint molecular methods when broken down by genotypic class: standard homozygotes, heterozygotes, and inverted homozygotes. While we have no objective measure of which specimens are ‘true’ heterozgyotes and ‘true’ inverted homozygotes, it is noteworthy that the discordances for all inversions other than 2Rc in *An. gambiae* either skew toward molecular genotypes of ‘1’ or ‘2’, or they are roughly equally distributed between ‘1’ or ‘2’ and ‘0’. The pattern for 2Rc in *An. gambiae* is distinctive, in that the skew is strongly toward molecular genotypes of ‘0’, which is consistent with tags that may not be appropriately suited for the *An. gambiae* population in which they are applied. Further study is both required and merited, to understand the cause(s) of population structure between the populations used to develop the 2Rc tags for in *An. gambiae* in Ag1000G and those used to test the tags in the present study. Interestingly, 2Rc contains cytochrome P450 genes implicated in insecticide resistance in *An. gambiae* and *An. coluzzii* (Love *et al.* 2016; Main *et al.* 2015), and more broadly, Coluzzi and colleagues (2002) observed that the region spanned by 2Rc is involved in many rearrangements that differentiate members of the *An. gambiae* species complex, leading these authors to propose that this region may have ecological relevance with respect to larval breeding site adaptations.

**Table 3.** Concordance between CYT and both molecular methods by inversion genotype *(0, 1, 2)* for specimens for which both molecular methods agree. Shown are the numbers of specimens scored for each pairwise comparison.

|  |  | GT-seq + OA |  |  |
| --- | --- | --- | --- | --- |
|  | CYT | 0 | 1 | 2 |
| **2La** | 0 | **1** | 0 | 0 |
|  | 1 | 0 | **80** | 2 |
|  | 2 | 1 | 70 | **794** |
| **2Rb** | 0 | **124** | 1 | 7 |
|  | 1 | 1 | **491** | 9 |
|  | 2 | 7 | 16 | **249** |
| **2Rd** | 0 | **389** | 7 | 0 |
|  | 1 | 7 | **31** | 0 |
|  | 2 | 0 | 0 | **0** |
| **2Ru** | 0 | **778** | 39 | 10 |
|  | 1 | 7 | **84** | 0 |
|  | 2 | 0 | 0 | **8** |
| **2Rc_col** | 0 | **153** | 7 | 0 |
|  | 1 | 10 | **235** | 0 |
|  | 2 | 0 | 1 | **95** |
| **2Rc_gam** | 0 | **309** | 1 | 0 |
|  | 1 | 34 | **43** | 0 |
|  | 2 | 4 | 0 | **0** |

CYT, cytogenetics; OA, Open Array

Although we find good concordance between both molecular genotyping approaches, GT-seq more often agreed with cytogenetics than did OA (Table 2). This is not surprising, given the hybridization-based nature of OA and the extremely high levels of nucleotide diversity found in *An. coluzzii* and *An. gambiae* (Miles *et al.* 2017). In addition, highly concordant candidate tag SNPs that also had low polymorphism in the ~50 bp immediately surrounding the tag, as required by OA, were sufficiently rare that we were compelled to lower the concordance threshold to find enough candidates suitable for assay design, and consequently the total number of OA tags per inversion is smaller than for GT-seq (Table 1). These factors likely compound to lower the performance of OA compared to GT-seq. Furthermore, as detailed by Campbell *et al.* (2015), genotyping costs are lower for GT-seq compared with OA. Nevertheless, if the number of tag SNPs to be genotyped is low (50-100) and the number of samples high (10^2^ to 10^3^), OA remains a cost effective option and is still widely used (Campbell *et al.* 2015).

## CONCLUSIONS

Chromosomal inversions have been viewed as instruments of ecotypic differentiation in anopheline mosquitoes (Coluzzi 1982). Insights into their adaptive significance as balanced polymorphisms and their possible role in behavioral variation, optimal habitat choice, and the speciation process (Coluzzi 1982) were gained from extensive polytene chromosome analyses largely conducted in the pre-genomic era (Coluzzi *et al.* 2002; Coluzzi *et al.* 1979; Manoukis *et al.* 2008; Toure *et al.* 1998). Now, with access to reference genome assemblies and powerful functional genomics tools, the potential exists to probe molecular mechanisms and deepen our understanding. Yet, a major limitation to progress in this area has been the strict requirement for polytene chromosome analysis, which not only limits samples but also demands rare cytogenetic expertise whose throughput is low. Here we demonstrate that tag SNPs highly correlated with inversion status can be used for joint molecular genotyping of common inversions in *An. gambiae* and *An. coluzzii* across the genome (*i.e.*, for karyotyping). Molecular genotyping methods, both OA and GT-seq, can be performed at scale and the results are comparable or superior to traditional cytogenetic karyotyping. These tools invite a renewal of investigations into the role of chromosomal inversions in anopheline behavior and environmental adaptation.

## ACKNOWLEDGEMENTS

We thank the technicians of IRD/IRSS/Centre Muraz of Burkina Faso for their fundamental contribution to the field work, as well as the inhabitants of the sampled villages for their kind collaboration. We thank M. Kern and R. Montañez-Gonzalez (University of Notre Dame) for assistance with DNA extraction. The Notre Dame Center for Research Computing provided technical support. We thank Dana-Farber/Harvard Cancer Center in Boston, MA, for the use of the Genotyping and Genetics for Population Sciences Core, which provided SNP analysis using the Taqman OpenArray Genotyping System. We acknowledge the Notre Dame Genomics and Bioinformatics Core Facility for assistance with GT-seq project development, library preparation, and sequencing. This work was supported by the National Institutes of Health (R01 AI125360 awarded to NJB). During this work, NJB was supported by Target Malaria, which receives core funding from the Bill & Melinda Gates Foundation and from the Open Philanthropy Project Fund, an advised fund of Silicon Valley Community Foundation. Dana-Farber/Harvard Cancer Center is supported in part by an NCI Cancer Center Support Grant # NIH 5 P30 CA06516.

